# ANALYSIS OF MICROBIAL DIVERSITY IN RAW FISH CEVICHE

**DOI:** 10.1101/2020.06.24.170217

**Authors:** M. E. Ramírez-Martínez, G. C. Rodríguez-Castillejos, M. C. Hernández-Jiménez, L. Y. Ramírez-Quintanilla, F. Siller-López, E. Acosta-Cruz, H. Martínez-Montoya

## Abstract

Ceviche is a traditional dish made from raw fish meat marinated in lime juice without any heat cooking step throughout its preparation process. Although the use of organic acids as antibacterial agents is well known; recent research indicates that lime juice actually can reduce the risk of *V. parahemolyticus* infections but it is ineffective against other potential pathogens. Despite the fact that fresh fish meat is safe; exposed organs including skin, gills and guts represent a potential source of bacterial contamination. In Mexico, diarrheal diseases are caused mainly by contaminated food; it is estimated that almost 67% of infections are due the presence of bacterial agents mainly in frozen and fresh fish. The main objective of this study was to estimate the taxonomic diversity of microbial species present in ready-to-eat ceviche using a metagenomic approach. Six samples of commercially available ceviche were subjected to DNA high throughput sequencing and bioinformatics analyses, we identified between 65,000 and 131,000 reads per sample. The predominant phyla identified through the samples were *Proteobacteria*, *Bacteroidetes* and *Firmicutes*. We discuss the factors involved in the microbiological quality of this kind of raw foods and how they influence the bacterial diversity within the analyzed samples.

## INTRODUCTION

Ceviche is a traditional dish in many Latin American countries but especially popular in Mexico and Peru. It is made using raw fish meat, marinated few hours in citrus juices in order to eliminate potential pathogens and add the meat a cooked-like flavor and consistency. Lime juice from key limes (*Citrus aurantifolia*) is one of the most widespread ingredients in its preparation despite the manufacturing region. However, depending of where it is made, other citrus juices such as sour orange and weak organic acids as acetic acid could be used. These products have been reported as effective antibacterial agents due their dissociation inside the bacterial cell into a proton that ultimately will reduce the cytoplasm pH and an anion that will induce metabolic perturbations in the acidified cytoplasm (Hirshfield, Terzulli et al. 2003). Lime juice has shown antibacterial activity against *Vibrio parahemolyticus* and *V. cholerae* when used in raw seafood (Tomotake, Koga et al. 2006) and it exhibited inactivation of common foodborne bacteria such as *E. coli* O157:H7, *S. enteriditis* and *L. monocytogenes* from marinated raw beef (Yang, Lee et al. 2013).

Bacterial hazards associated to seafood consumption are divided in two groups: the first composed by indigenous bacteria naturally present in the aquatic environment; the second group are introduced bacteria as a result of human, animal or otherwise contamination in the aquatic environment or during the seafood processing and handling. Although ingredients and fish meat used for ceviche may vary by region, in Mexico it is made with either marine fish such as tuna (*Thunnus* sp.), mullet (*Mugil cephalus*), sea brass (*Dicentrachus labrax*) and freshwater species including catfish (*Ictalurus punctatus*), seabream (*Diplodus vulgaris*), trout (*Oreochromis mykiss*) and tilapia (*O. niloticus*). Fresh fish meat is known to be free of pathogens, however, exposed body structures like skin, gills and guts could increase the risk of foodborne diseases due to parasitic, viral or bacterial contamination. Moreover, poor hygienic handling, transportation and preparation processes could also increase the risk of pathogenic bacteria occurrence and ultimately alter significantly the structure of microbial communities present in seafood. In order to detect such pathogens, traditionally, food microbiology relies on culture-based techniques. However, in addition to their low sensitivity, those techniques need growth conditions that are usually difficult to reproduce in the laboratory, and this could lead to underestimations or bias during the determination of microbial diversity. On the other hand, culture-independent methods based in the sequencing of nucleic acids purified directly from the environmental sample, could improve significantly the limitations exhibited by the classic microbiological methods (De Filippis, Parente et al. 2018). In the present study, we analyzed six ceviche samples through massive sequencing of the 16S V3-V4 rRNA amplicon to estimate the bacterial diversity present in this kind of raw seafood. We obtained between 65,000 and 131,000 reads within the sequenced samples, the reads were quality filtered and assembled in contigs. The resulting contigs were compared and identified by homology using the small ribosomal subunit (SSU) SILVA database v. 132. The predominant phyla found within the samples were *Bacteroidetes*, *Proteobacteria*, *Firmicutes* and *Actinobacteria* while at genus level, *Psychrobacter*, *Shewanella*, *Prevotella*, *Pseudomonas* and *Pantoea* were found the most abundant.

## MATERIAL AND METHODS

### Sample collection

Ceviche samples, identified as S12, S13, S14 and S15 were obtained from restaurants and street vendors in Reynosa, Mexico. Each sample was transported on ice to the laboratory in sterile plastic bags and stored at −20 C before DNA purification and further analysis. Additionally, samples S16 and S17 were prepared in the laboratory from 100 g of chopped mullet and seabream respectively and were marinated with 8 mL of lime juice two hours before processing.

### DNA purification and sequencing

Ceviche samples were homogenized in 0.9% sterile saline solution before DNA purification. Total genomic DNA was recovered from 0.25 grams of sample using a modified chloroform-ethanol procedure using 1 mL of lysis buffer (0.5M EDTA, 10% SDS) and precipitated in two steps with isopropanol and ethanol. Finally, it was diluted in 50 μL of AE buffer (Qiagen, Valencia, CA). Prior the library preparation, DNA quality control was performed with Bioanalyzer (Agilent, Santa Clara, CA) to determine the size of DNA fragments and its abundance throughout the samples. DNA pair-ended library was performed according to Illumina’s Multiplexing Sample Preparation Guide (Illumina, San Diego, CA), the DNA was fragmented, end repaired, A’ tagged, ligated to 357-F and CD-R adaptors, size-selected, and enriched with 25 cycles of PCR during which an index was incorporated to the samples (Table S1). The resulting libraries were subjected to Illumina MiSeq (2×300) pair-ended sequencing at the Laboratory of Genomic Services of the National Laboratory of Genomics for Biodiversity (LANGEBIO) (Irapuato, Mexico).

### Sequencing data analyses

Taxonomic labels were assigned to the raw reads in each sample by homology with a local 16S rRNA database using Kraken 2 (Wood and Salzberg 2014) in the Illumina BaseSpace Sequence Hub (https://basespace.illumina.com). Additionally, sequencing reads were assembled in contigs and quality filtered following the Mothur pipeline (Schloss, Westcott et al. 2009) to assess species richness and alpha diversity.

Contigs were compared and identified by homology with the ribosomal small subunit (SSU) SILVA Database v. 132 (Quast, Pruesse et al. 2013). Aligned sequences were screened and to non-bacterial and chimeric sequences were removed. The alpha diversity for each sample was reported as observed and calculated using the Chao1, ACE and Simpson’s diversity metrics and visualized throughout the Microbiome Analyst Portal (https://www.microbiomeanalyst.ca) (Dhariwal, Chong et al. 2017). Finally, metagenomeSeq (Paulson, Stine et al. 2013) was used to find differential abundance of microorganisms at different taxonomic levels among commercial and control samples.

## RESULTS

Sequenced reads were deposited and are available in the NCBI sequence depository under the Raw Fish Ceviche Bacterial Diversity BioProject (ID: PRJNA512206). We identified between 65,000 and 131,000 reads among the sequenced samples. According to the taxonomic labels given by both pipelines, Kraken and Mothur, the predominant phyla across the ceviche samples were *Proteobacteria*, *Actinobacteria*, *Firmicutes* and *Bacteroidetes*. Additionally, we found several genera with low count of reads, *Chloroflexi*, *Cyanobacteria*, *Fusobacteria* and *Gemmatimonadetes*. At genus level, Kraken analysis showed high predominance of *Bacteroidetes* followed by *Firmicutes* in S12 but S13 had high presence of *Firmicutes*, whereas S14, S15, S16 and S17 showed predominance of bacteria belonging to the phylum Proteobacteria (Table 1).

**Table 1.**
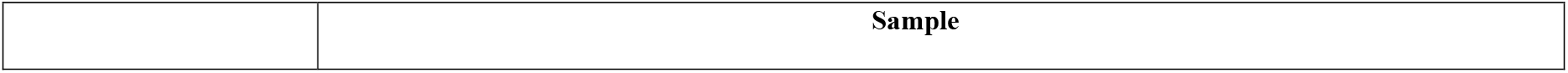
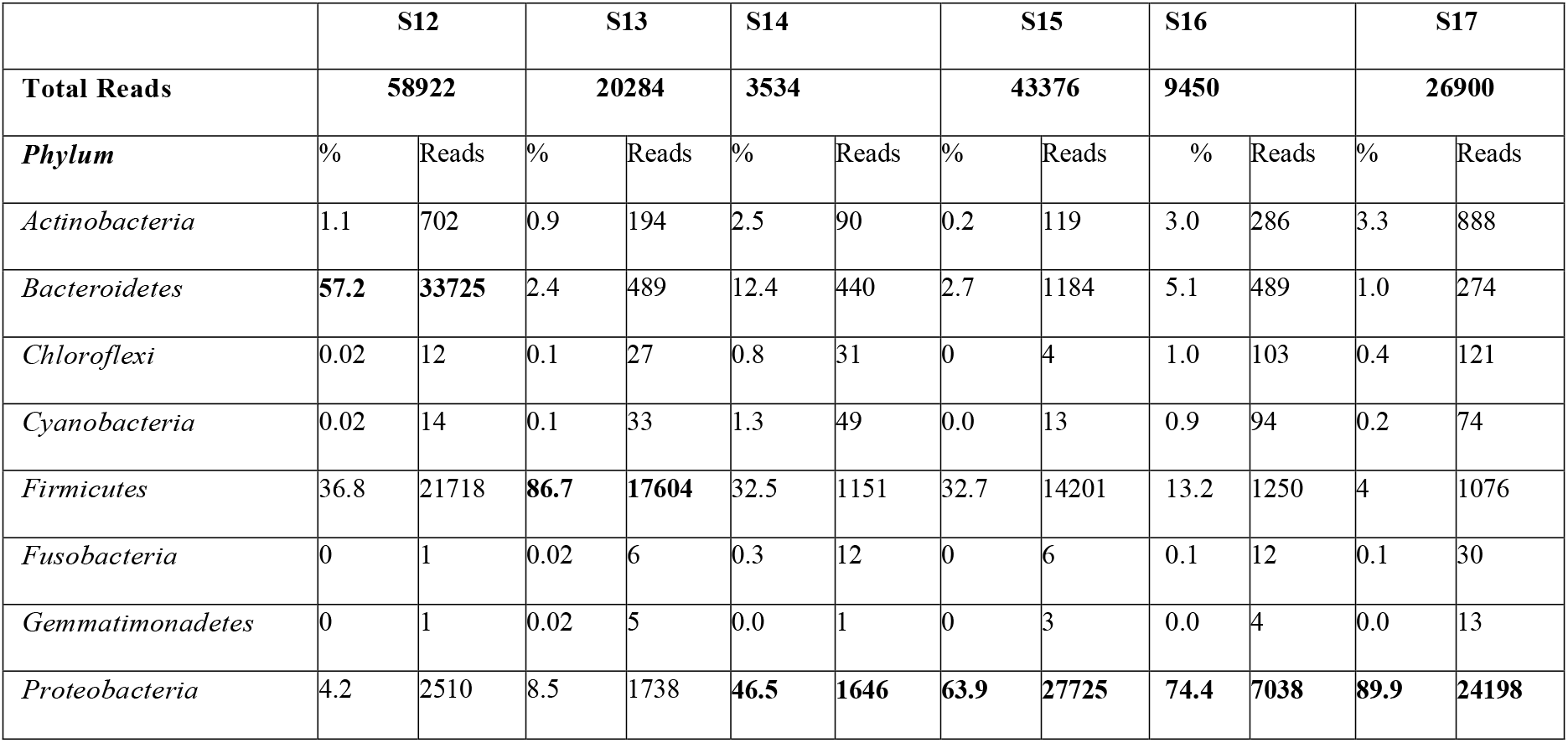
Total reads identified to phylum for each sample tested in Kraken (numbers in bold represent the most representative phylum within the sample)

|  | S12 |  | S13 |  | S14 |  | S15 |  | S16 |  | S17 |  |
| --- | --- | --- | --- | --- | --- | --- | --- | --- | --- | --- | --- | --- |
| Total Reads | 58922 |  | 20284 |  | 3534 |  | 43376 |  | 9450 |  | 26900 |  |
| Phylum | % | Reads | % | Reads | % | Reads | % | Reads | % | Reads | % | Reads |
| *Actinobacteria* | 1.1 | 702 | 0.9 | 194 | 2.5 | 90 | 0.2 | 119 | 3.0 | 286 | 3.3 | 888 |
| *Bacteroidetes* | 57.2 | 33725 | 2.4 | 489 | 12.4 | 440 | 2.7 | 1184 | 5.1 | 489 | 1.0 | 274 |
| *Chloroflexi* | 0.02 | 12 | 0.1 | 27 | 0.8 | 31 | 0 | 4 | 1.0 | 103 | 0.4 | 121 |
| *Cyanobacteria* | 0.02 | 14 | 0.1 | 33 | 1.3 | 49 | 0.0 | 13 | 0.9 | 94 | 0.2 | 74 |
| *Firmicutes* | 36.8 | 21718 | 86.7 | 17604 | 32.5 | 1151 | 32.7 | 14201 | 13.2 | 1250 | 4 | 1076 |
| *Fusobacteria* | 0 | 1 | 0.02 | 6 | 0.3 | 12 | 0 | 6 | 0.1 | 12 | 0.1 | 30 |
| *Gemmatimonadetes* | 0 | 1 | 0.02 | 5 | 0.0 | 1 | 0 | 3 | 0.0 | 4 | 0.0 | 13 |
| *Proteobacteria* | 4.2 | 2510 | 8.5 | 1738 | 46.5 | 1646 | 63.9 | 27725 | 74.4 | 7038 | 89.9 | 24198 |

At genus level, in S12 we have found a high prevalence of *Bacteroidetes*, mainly represented by *Prevotella* species (58%) such as *P. oris*, *P. dentalis*, *P. fusca*, *P. denticola*; the second largest group of bacteria in S12 belongs to the phylum *Firmicutes* and had homologous sequences to species such as *Faecalibacterium prausnitzii*, *Eubacterium rectale*, *Ruminococcus bicirculans*, *Selenomonas sputigena* and *S. ruminantium*, *Megaspharea elsdenii* and in less frequency, we found *S. aureus* and *Streptococcus* sp. In S13 we observed a high relative abundance (RA) of not assigned bacterial genus (72.2%) but also we found *Psychrobacter* (9.34%), *Rothia* (5.08%) and *Paracoccus* (3.7%) additionally represented in less than 2% of RA, *Enterobacteria*, *Pantoea*, *Propionibacterium* and *Shewanella*. Sample S15 had 0.95% of RA of *Empedobacter* and *Rheinheimera*. S14 had a high presence of *Pantoea* (50%) and genera such as *Rosenbergiella* (16.95%), unclassified *Enterobacteriaceae* (11.96%) and *Weisella* (5.46%). In less percentage, we found *Myroides*, *Pseudomonas*, *Chryseobacterium*, *Rhizobium*, *Buttiauxella*, *Sphingobacterium*, *Soonwooa*, *Rothia* and *Pseudochrobacterium*. Non-commercial samples, S16 and S17 were only fish meat marinated with lime juice. Whereas S16 exhibited *Psychrobacter* (16.45%), *Pseudomonas* (3.5%), *Propionibacterium* (1.9%), *Shewanella*, *Roseiflexus*, *Flavobacterium*, *Acinetobacter*, *Rothia*, *Bradirhizobium* and *Aeromonas* on the other hand, S17 showed *Aeromonas* (19.13%), *Shewanella* (8.77%), *Propionibacterium* (8.09%), *Psychrobacter* (3.8%) and less represented, *Corynebacterium*, *Acinetobacter*, *Roseiflexus*, *Pseudomonas*, *Flavobacterium* and *Buttiauxella*.

Accordingly, to the Kraken analysis, Mothur analyses revealed a high abundance of *Bacteroidetes* in the sample S12 but we found a relatively low abundance of *Firmicutes*, and the most abundant phyla in S13, S14, S15, S16 and S17 were *Proteobacteria*, *Actinobacteria*, *Firmicutes*, *Chloroflexi* and *Armatimonadetes* (Figure 1 and Table S2).

**Fig 1.**
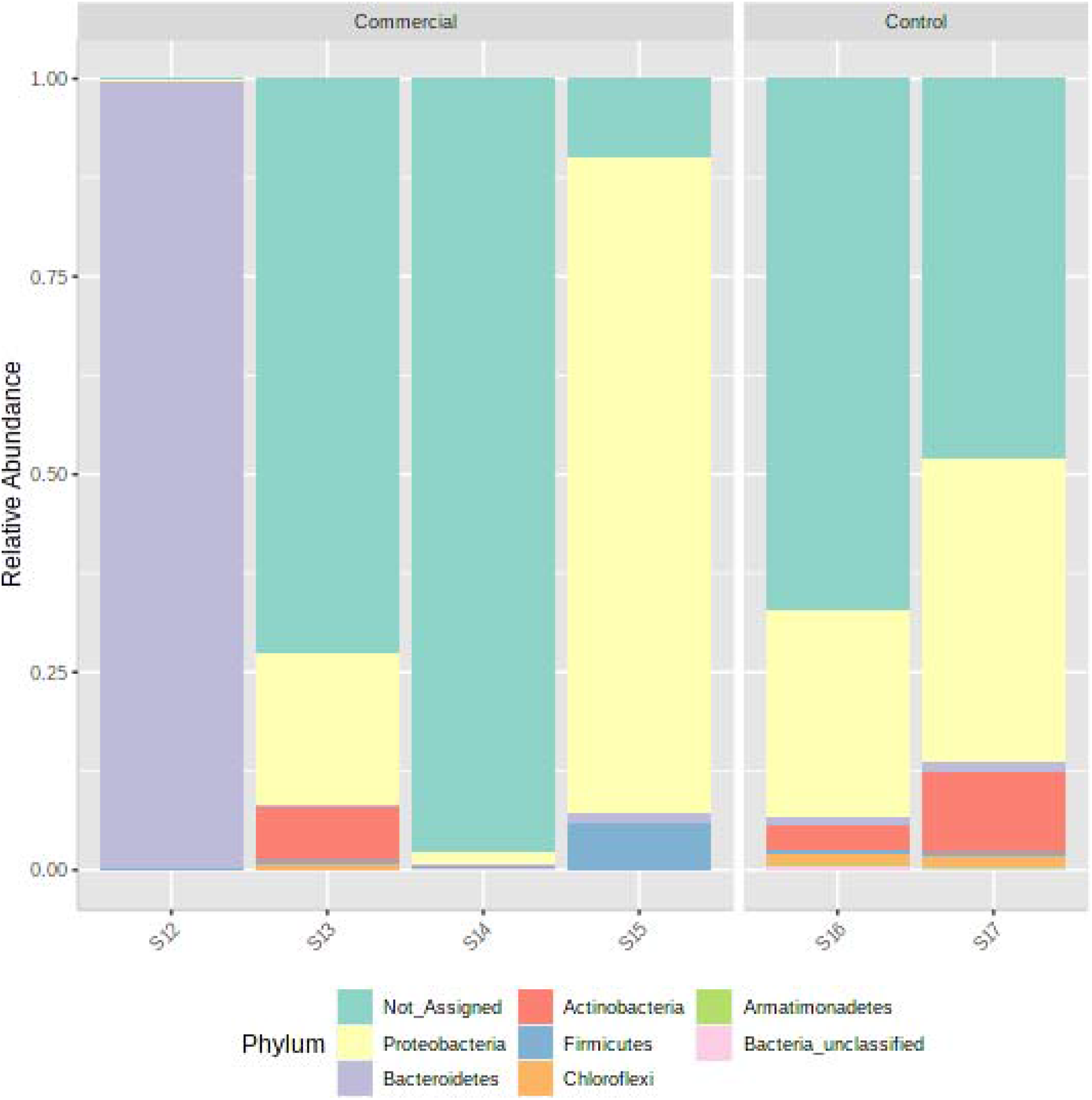
Dominant phyla across analyzed ceviche samples.

At genus level, we found mostly *Prevotella* in S12 (0.995) while in S13-S15 a higher diversity was observed (Figure 2). The rarefaction richness plot observed in samples was not saturated in the ceviche samples but it clearly indicated that sample S12 had the lowest richness while S17 showed the highest number of OTU’s. (Figure 3 and Table S3).

**Figure 2.**
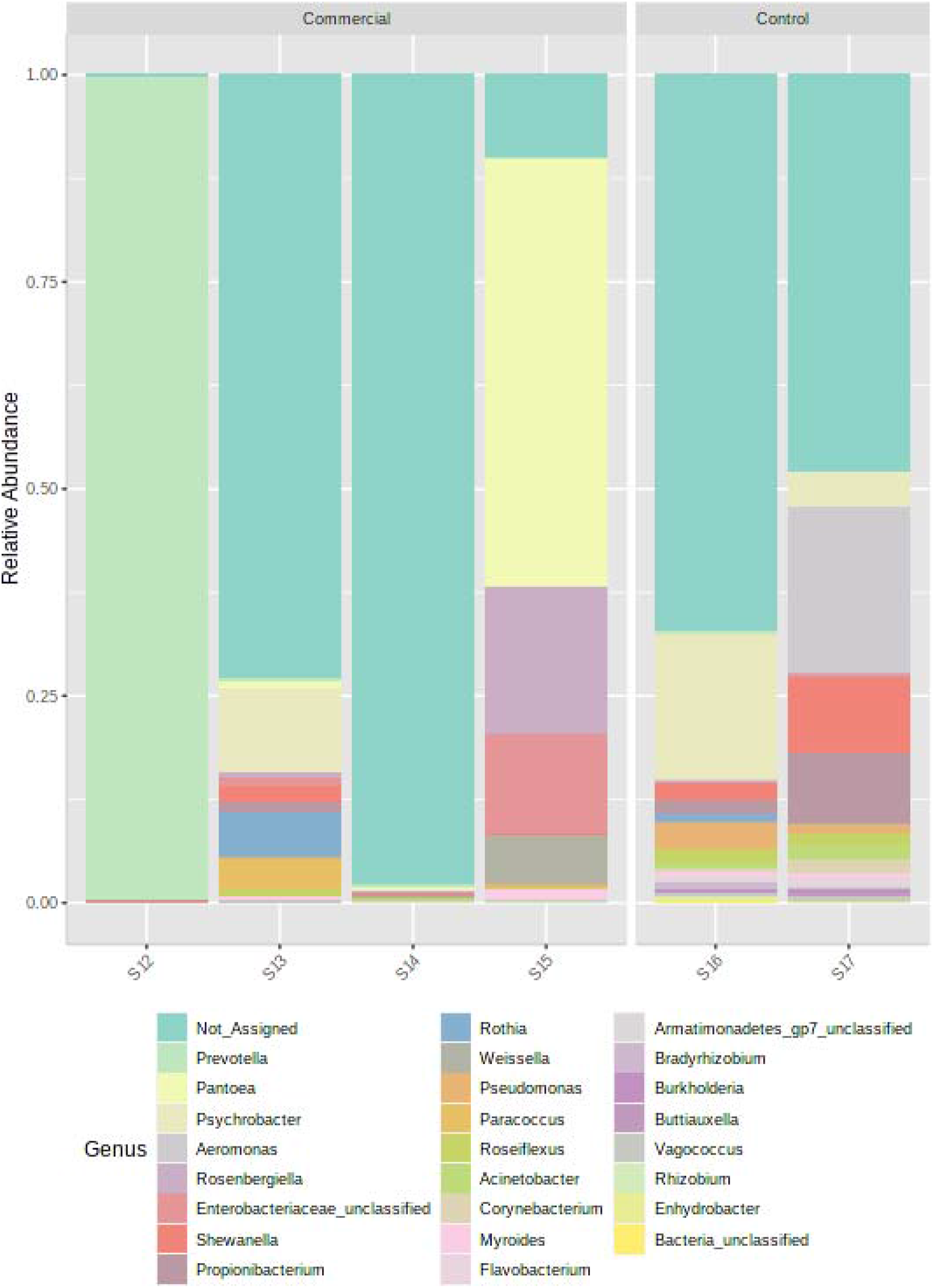
Relative abundance of bacterial genera within the Ceviche samples.

**Figure 3.**
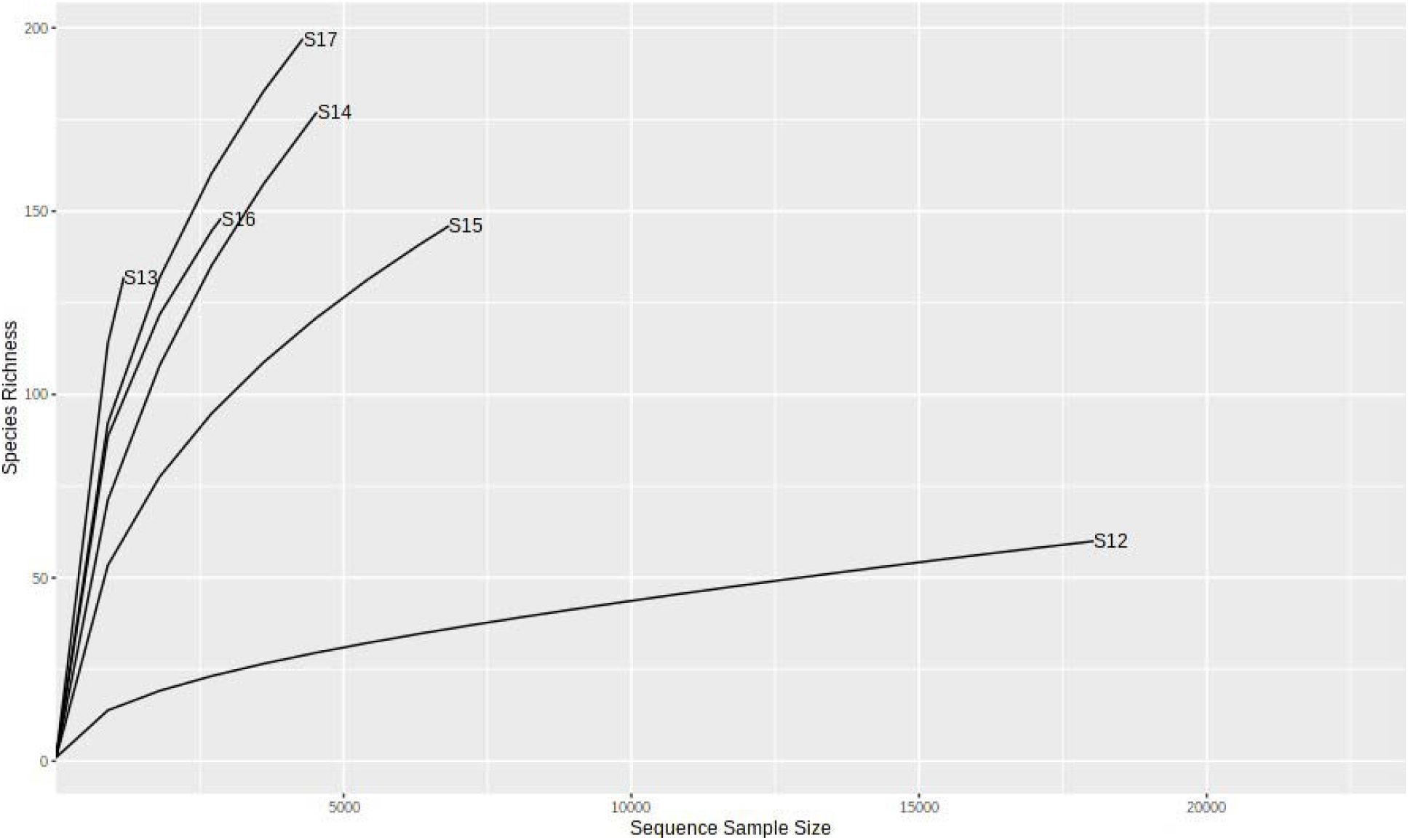
Richness rarefaction plot of analyzed ceviche samples.

Alpha diversity was assessed through observed and estimated analyses using the Mothur data. We did not found significant differences in observed diversity among samples (*P*=0.29724). however, according to the rarefaction analysis, S12 and S17 had the lowest and highest diversity with 60 and 197 OTU’s respectively (Table 2). In addition to the observed count of species we calculated the alpha diversity trough ACE (*P*= 0.71295), Chao1 (*P*=0.98386) and Simpson’s diversity indexes (*P*=0.2646) (Table 2).

**Table 2.** Alpha diversity observed and calculated for analyzed samples.

| Sample type | Sample ID | Chao1 |  | ACE |  | Simpson |  | Observed |  |
| --- | --- | --- | --- | --- | --- | --- | --- | --- | --- |
|  |  | Value | SE | Value | SE | Value | SE | Value | SE |
| Commercial | S12 | 8.33 | 0.91 | 11.27 | 1.79 | 0.009 | NA | 60 | NA |
| Commercial | S13 | 24.75 | 4.19 | 27.82 | 2.71 | 0.45 | NA | 132 | NA |
| Commercial | S14 | 17.2 | 0.61 | 17.94 | 1.85 | 0.04 | NA | 177 | NA |
| Commercial | S15 | 18.5 | 2.57 | 19.6 | 1.9 | 0.67 | NA | 146 | NA |
| Control | S16 | 23 | 0.48 | 23.26 | 2.31 | 0.51 | NA | 148 | NA |
| Control | S17 | 22 | 0.48 | 22.38 | 1.92 | 0.71 | NA | 197 | NA |

Correlation and clustering visualization analysis showed that based on the microbial profiles, laboratory-made samples S16 and S17 were clustered together whereas commercial samples formed the other cluster. It is observed that S16 and S17 samples were strongly positively correlated to the presence of bacteria belonging to the *Psychrobacter*, *Pseudomonas*, *Bradyrhizobium*, *Shewanella*, *Propionibacterium*, *Corynebacterium*, *Aeromonas*, *Flavobacterium*, *Acinetobacter* and *Vagococcus* genera which do not suggest evidence of human contamination (Figure 4). On the other hand, S12, S13 and S14 had a strong correlation with bacterial classes typically associated to human contamination such as *Enterobacteriales Lactobacillales*, *Bacteroidales* and *Rhodobacterales*. We found that *Enterobacteriaceae* (*P*= 2.63E-4), *Prevotellaceae* (*P*= 0.005) and *Leuconostocaceae* (*P*= 0.005) family members had a significant higher abundance in ceviche samples from restaurants than in the laboratory made samples whereas *Moraxellaceae* was more slightly abundant in the control samples (*P*= 0.02) (Table S4).

**Figure 4.**
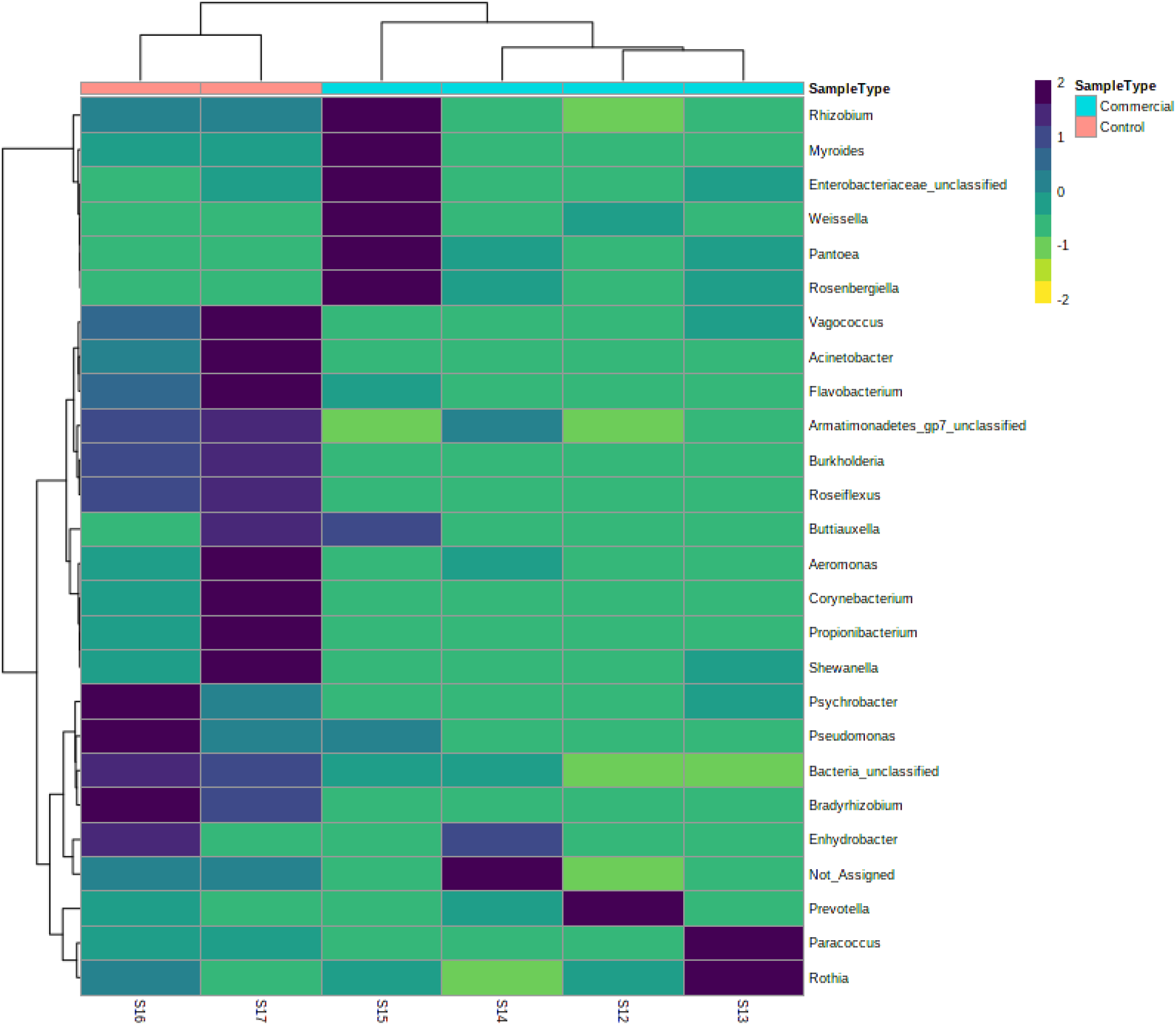
Hierarchical clustering and heatmap visualization of commercial and laboratory made samples at genus level.

## DISCUSSION

Bacterial agents found in the tested ceviche samples showed high diversity of taxa across the samples, as stated previously. In general, bacterial in seafood belong to two different groups, native marine or freshwater bacteria present in naturally in the aquatic environment and those bacteria present due exogenous contamination of human or animal sources.

Among the native bacteria identified in this study we found *Soonwooa*, a member of the *Flavobacteriaceae* family is a previously isolated microorganism from the yellow sea waters from Korea (Joung, Song et al. 2010), it was found mainly in S14 and S15. Nowadays, two species of *Soonwooa*, *S.* sp. and *S. buanensis* have been described in tissues from tilapia (*O. niloticus*) reared in dams in Mexico (Soto-Rodriguez, Cabanillas-Ramos et al. 2013) and in duck meat products from Korea (Kim, H.I. et al. 2016) but were not identified as pathogenic agents. Within the *Flavobacteriaceae* family we also identified reads hitting to the gram-negative *Chryseobacterium* in S13 and S15, although we were not able to identify the bacteria at species level, its close relatives, previous reports mention that *C. meningosepticum* has been associated to foodborne septicemia due consumption of raw seafood (Kim, H.I. et al. 2016) and *C. indologenes* which is widely spread in soil and plants rarely was associated to human disease (Imataki and Uemura 2017). This last example could be present in ceviche since the samples involved raw vegetables in its preparation, however, further analyses are required to elucidate the origin of *Chryseobacterium* species present in raw seafood.

The genus *Rheinheimera*, found in S14 is widely distributed in freshwater, saltwater and soil environments. Is a Gram-negative bacterium that presents antimicrobial activity in some strains (Chen, Lin et al. 2010). Additionally, other bacteria naturally associated to aquatic environments such as *Shewanella*, *Psychrobacter*, *Vagococcus*, *Buttiauxella* and *Novosphingobium* were found, although they do not represent a potential source of foodborne illness Vegetables in ceviche, including cilantro, onion, tomato, pepper and avocado are usually consumed raw; in ceviche, they represent a major source of pathogenic bacteria that is associated to foodborne illness. In this study the ceviche samples tested had a variety of produce ingredients that constituted a factor of variability in the results. Among exogenous bacteria, we identified several members of the *Enterobacteriaceae* genus such as *Pantoea*, *Rosenbergiella* and *Morganella*. *Pantoea*, primarly found in S15 but also in S13, these genera are known to be associated as a pathogen or epiphytic bacteria in edible vegetables, e.g *P. agglomerans* in tomato and onion (Oie, Kiyonaga et al. 2008, Nadarasah and Stavrinides 2014) which are main ingredients of Mexican made ceviche. *P. agglomerans*, has also been associated to soft bone infections [41], although this opportunistic bacterium is not known to be a foodborne pathogen it has been associated as a contaminant in powered infant formula milk (Mardaneh and Dallal 2013). We hypothesized that *Pantoea* sp. found in ceviche have its origin in the vegetables used in the dish preparation since it was almost undetectable in the laboratory made samples S16 and S17 respectively.

*Rothia* sp., found in a highest frequency in S13 may cause a significant number of infections to immunocompromised hosts despite they are part of the normal microbiome of human oropharynx and respiratory tract (Ramanan, Barreto et al. 2014), it is possible that *Rothia* in ceviche have its origin from anthropogenic contamination during its preparation.

The genus *Weissella*, found primarily in S15 but also in S13 is a gram-positive cocco-bacilli, often misidentified by traditional microbiological methods but commonly found in fermented foods, including vegetables, meat, fish and raw milk. Although, this genus is rarely a known cause of human infections. Conversely, few *Weissella* strains may act as probiotics due its antimicrobial activity e.g. *W. hellenica* isolated from flounder intestine act as a defensive agent against host pathogens such as *Edwardsiella*, *Pasteurella*, *Aeromonas* and *Vibrio* (Cai, Benno et al. 1998, Abriouel, Lerma et al. 2015).

*Aeromonas*, a genus found in high presence in S17, but present in all the tested samples is one of the etiological agents associated to disease-causing pathogen of fish and responsible of infectious complications in humans. This genus can be isolated from almost every ecological niche in earth, including aquatic habitats, fish, food and soils and it can be present in ceviche due multiple factors and represented by one or more species. In food, *Aeromonas* strains have been widely reported in dairy products, specifically unpasteurized milk, yogurt and cheese from diverse geographical origins.

*Prevotella*, was detected in all samples except S15 and S17, this agent is known to be a part of human gut, oral and vaginal microbiota, this bacterium is often an indicator of fecal contamination of water (Fogarty and Voytek 2005, Koskey, Fisher et al. 2014). However, in ceviche the high presence of this agent could be associated to low hygienic conditions in produce handling.

*Pseudomonas*, a naturally widespread genus is a potential human pathogen previously reported in spinach, lettuce and red cabbage, it was found in every sample and probably its presence is due either contaminated vegetables and contamination of fish meat. In the present study, we did not found etiological agents associated to freshwater fish species outbreaks of foodborne illness such as *Salmonella*, *Escherichia coli*, *Clostridium*, *Staphylococcus* or *Bacillus*, this study presents the first description of the bacterial diversity in raw fish food using a metagenomic approach. However, as observed, microbiological quality of raw foods, depends on many factors bacterial diversity in raw fish ceviche increases as raw vegetables are added. Further analyses are needed in order to examine the dynamics of bacterial communities in ready-to-eat raw seafood.

## Acknowledgments

Authors thank to the Secretaría de Educación Pública of Mexico for the funding given to this study through the PRODEP-NPTC fellowship granted to Humberto Martinez Montoya and Myriam Elizabeth Ramirez Martinez (UAT-PTC-211)

