## Supplemental Table 1 for "ANALYSIS OF MICROBIAL DIVERSITY IN RAW FISH CEVICHE": Supplementary Table 1.docx

**Table S1.** Indexes and identified reads of sequenced libraries

| **Sample** | **Library** | **Index 1 (i7)** | **Index 2 (i5)** | **Identified reads** |
| --- | --- | --- | --- | --- |
| S12 | HM1CM1SS09 | GGAGCTAC | GCGTAAGA | 130,147 |
| S13 | HM1CM1SS10 | GGAGCTAC | CTATTAAG | 109,459 |
| S14 | HM1CM1SS11 | GGAGCTAC | AAGGCTAT | 108,256 |
| S15 | HM1CM1SS12 | GGAGCTAC | GAGCCTTA | 131,305 |
| S16 | HM1CM1SS13 | GGAGCTAC | TTATGCGA | 65,259 |
| S17 | HM1CM1SS14 | GCGTAGTA | TGCACTAG | 94,001 |
