## Supplemental Table 2 for "ANALYSIS OF MICROBIAL DIVERSITY IN RAW FISH CEVICHE": Supplementary Table 2.docx

**Table S2.** Relative abundance of phyla

|  | **S12** | **S13** | **S14** | **S15** | **S16** | **S17** |
| --- | --- | --- | --- | --- | --- | --- |
| *Actinobacteria* | 0.001 | 0.068 | 0.004 | 0.003 | 0.033 | 0.104 |
| *Armatimonadetes* | 0.000 | 0.002 | 0.002 | 0.000 | 0.005 | 0.005 |
| *Bacteria_unclassified* | 0.000 | 0.000 | 0.002 | 0.000 | 0.004 | 0.003 |
| *Bacteroidetes* | 0.995 | 0.004 | 0.002 | 0.013 | 0.011 | 0.011 |
| *Chloroflexi* | 0.000 | 0.007 | 0.001 | 0.000 | 0.014 | 0.013 |
| *Firmicutes* | 0.001 | 0.003 | 0.000 | 0.057 | 0.002 | 0.003 |
| *Not_Assigned* | 0.003 | 0.728 | 0.978 | 0.101 | 0.672 | 0.480 |
| *Proteobacteria* | 0.000 | 0.188 | 0.013 | 0.826 | 0.259 | 0.382 |
