## Supplemental Table 3 for "ANALYSIS OF MICROBIAL DIVERSITY IN RAW FISH CEVICHE": Supplementary Table 3.docx

**Table S3.** Relative abundance of bacterial genus

| **Genus** | **S12** | **S13** | **S14** | **S15** | **S16** | **S17** |
| --- | --- | --- | --- | --- | --- | --- |
| *Acinetobacter* | 0 | 0.0019 | 0 | 0 | 0.0095 | 0.0181 |
| *Aeromonas* | 0 | 0.0038 | 0.0028 | 0.00093 | 0.0049 | 0.2015 |
| *Armatimonadetes_gp7_unclassified* | 0 | 0.0019 | 0.0016 | 0 | 0.0049 | 0.0045 |
| *Bacteria_unclassified* | 0 | 0 | 0.00071 | 0.00031 | 0.0030 | 0.0015 |
| *Bradyrhizobium* | 0 | 0 | 0 | 0 | 0.0076 | 0.0033 |
| *Burkholderia* | 0 | 0.0009 | 0.0004 | 0 | 0.0038 | 0.0035 |
| *Buttiauxella* | 0 | 0 | 0 | 0.0026 | 0 | 0.0056 |
| *Corynebacterium* | 6.73E-05 | 9.63E-04 | 0 | 1.56E-04 | 4.59E-03 | 1.76E-02 |
| *Enhydrobacter* | 0 | 0 | 0.00191 | 0 | 0.0042 | 0.0002 |
| *Enterobacteriaceae_unclassified* | 0 | 0.013 | 0.00047 | 0.122 | 0.0011 | 0.0061 |
| *Flavobacterium* | 0 | 0 | 0 | 0.00062 | 0.0057 | 0.0084 |
| *Myroides* | 0 | 0.0009 | 0.00047 | 0.0124 | 0.0022 | 0.003 |
| Not_Assigned | 0.0028 | 0.728 | 0.9787 | 0.101 | 0.673 | 0.48 |
| *Pantoea* | 0 | 0.0115 | 0.0038 | 0.516 | 0.0003 | 0 |
| *Paracoccus* | 0 | 0.0375 | 0.0002 | 0 | 0.0022 | 0.0015 |
| *Prevotella* | 0.995 | 0.0028 | 0.0011 | 0 | 0.0034 | 0 |
| *Propionibacterium* | 0.00013 | 0.0144 | 0.0038 | 0.00077 | 0.0206 | 0.0853 |
| *Pseudomonas* | 6.73E-05 | 9.63E-04 | 9.57E-04 | 5.30E-03 | 3.10E-02 | 9.96E-03 |
| *Psychrobacter* | 0.0001 | 0.097 | 0.00047 | 0.00015 | 0.1724 | 0.0401 |
| *Rhizobium* | 0 | 0.0009 | 0.00023 | 0.002 | 0.0022 | 0.0012 |
| *Roseiflexus* | 0 | 0.0067 | 0.00071 | 0 | 0.0137 | 0.013 |
| *Rosenbergiella* | 0 | 0.0028 | 0.00119 | 0.175 | 0 | 0 |
| *Rothia* | 0.0008 | 0.0529 | 0 | 0.002 | 0.008 | 0.0012 |
| *Shewanella* | 0 | 0.0163 | 0 | 0 | 0.018 | 0.0901 |
| *Vagococcus* | 0 | 0.0019 | 0 | 0.00015 | 0.0019 | 0.0028 |
| *Weissella* | 0.0006 | 0.00096 | 0 | 0.05643 | 0 | 0 |
