## Supplemental Table 4 for "ANALYSIS OF MICROBIAL DIVERSITY IN RAW FISH CEVICHE": Supplementary Table 4.docx

**Table S4.** Metagenomeseq analysis for bacterial genus differentialy abundant among commercial and laboratory made ceviche samples (* denotes significant difference among tested groups)

| **Bacterial genus** | ***P* values** | **FDR** |
| --- | --- | --- |
| *Pantoea ** | 2.78E-06 | 7.23E-05 |
| *Prevotella ** | 1.18E-05 | 0.0001537 |
| *Rosenbergiella ** | 2.30E-05 | 0.0001993 |
| *Enterobacteriaceae* unclassified * | 0.00022369 | 0.001454 |
| *Weissella ** | 0.0009371 | 0.0048729 |
| *Myroides ** | 0.011237 | 0.048696 |
| *Psychrobacter ** | 0.020585 | 0.076459 |
| *Shewanella* | 0.10497 | 0.34114 |
| *Buttiauxella* | 0.18803 | 0.48179 |
| *Not Assigned* | 0.20648 | 0.48179 |
| *Rhizobium* | 0.21417 | 0.48179 |
| *Paracoccus* | 0.22236 | 0.48179 |
| *Pseudomonas* | 0.28878 | 0.57757 |
| *Roseiflexus* | 0.47659 | 0.7998 |
| *Bradyrhizobium* | 0.50373 | 0.7998 |
| *Corynebacterium* | 0.53404 | 0.7998 |
| *Acinetobacter* | 0.55048 | 0.7998 |
| *Aeromonas* | 0.55371 | 0.7998 |
| *Enhydrobacter* | 0.61074 | 0.83575 |
| *Flavobacterium* | 0.66287 | 0.86173 |
| *Propionibacterium* | 0.73556 | 0.91069 |
| *Vagococcus* | 0.78716 | 0.93028 |
| *Rothia* | 0.842 | 0.94151 |
| *Armatimonadetes* unclassified | 0.89381 | 0.94151 |
| *Bacteria_unclassified* | 0.93359 | 0.94151 |
| *Burkholderia* | 0.94151 | 0.94151 |
